# The *HAC1* Histone Acetyltransferase Promotes Leaf Senescence via Regulation of *ERF022*

**DOI:** 10.1101/599902

**Authors:** Will E. Hinckley, Keykhosrow Keymanesh, Jaime A. Cordova, Judy A. Brusslan

## Abstract

Nutrient remobilization during leaf senescence nourishes the growing plant. Understanding the regulation of this process is essential for reducing our dependence on nitrogen fertilizers and increasing agricultural sustainability. Our lab is interested in chromatin changes that accompany the transition to leaf senescence. Previously, darker green leaves were reported for *Arabidopsis thaliana hac1* mutants, defective in a gene encoding a histone acetyltransferase in the CREB-binding protein family. Here, we show that two *Arabidopsis hac1* alleles display delayed age-related developmental senescence, but have normal dark-induced senescence. Using a combination of ChIP-seq for H3K9ac and RNA-seq for gene expression, we identified 44 potential HAC1 targets during age-related developmental senescence. Genetic analysis demonstrated that one of these potential targets, *ERF022*, is a positive regulator of leaf senescence. *ERF022* is regulated additively by HAC1 and MED25, suggesting MED25 recruits HAC1 to the *ERF022* promoter to increase its expression in older leaves.

## Introduction

Plants continuously produce new organs. During vegetative growth, new leaves form from the shoot apical meristem, and develop into protein-rich photosynthetic factories that export sugars. Eventually, the older leaves enter senescence by catabolizing the photosynthetic apparatus and exporting nitrogen-rich amino acids to support continuing growth (Himelblau and Amasino, 2001). Understanding the regulation of leaf senescence could maximize nitrogen recycling thus producing more nutrient-rich seeds and reducing the need for fertilizers.

The transition into leaf senescence is preceded (Kim et al., 2018a) and accompanied by changes in gene expression (Buchanan-Wollaston et al., 2005; van der Graaff et al., 2006; Breeze et al., 2011). Lists of senescence-associated genes (SAG) have been generated from these transcriptome analyses. Enriched biological processes from Gene Ontology (GO) analyses include response to the hormones salicylic acid (SA), jasmonic acid (JA), abscisic acid (ABA) and ethylene. Also, enrichment of GO terms autophagy, immune response, defense response, and response to reactive oxygen species demonstrates a molecular relationship between defense and leaf senescence. Additional GO terms highly represented in SAGs from age-related developmental senescence include response to chitin and glucosinolate biosynthesis (Brusslan et al., 2015). The consistent enrichment of the phosphorylation term among SAG lists is likely a result of high expression of receptor-like kinase gene-family members, which also are known to regulate defense (Antolín-Llovera et al., 2014).

Changes in chromatin structure are hypothesized to promote and/or maintain leaf senescence (Humbeck, 2013). We have previously shown a correlation between histone 3, lysine 4, trimethylation (H3K4me3) and histone 3, lysine 9 acetylation (H3K9ac) histone modifications and increased expression of senescence up-regulated genes (SURGs). A similar correlation was seen between histone 3, lysine 27 trimethylation (H3K27me3) marks and decreased expression of senescence down-regulated genes (SDRGs) (Brusslan et al., 2012; Brusslan et al., 2015). Genetic analysis suggests histone deacetylases regulate leaf senescence. HDA19 is a negative regulator of senescence (Tian and Chen, 2001) while HDA6 is a positive regulator of leaf senescence (Wu et al., 2008). HDA9 works with POWERDRESS to reduce the expression of four putative negative regulators of leaf senescence *(NPX1, TMAC2, WRKY57* and *APG9)*, thus promoting leaf senescence (Chen et al., 2016).

Recently, two studies linked chromatin changes to leaf senescence. The Polycomb Repressive Complex 2 (PRC2) catalyzes H3K27me3 for long-term repression of ABA-induced SAGs (Liu et al., 2018). Double mutants in two PRC2 subunits *(clf/swn)* retain high SAG expression even after these genes are repressed in WT. H3K27me3-target genes that continue to be expressed in *clf/swn* mutants are significantly enriched for leaf senescence-related GO terms, indicating that long-term dampening of SAG expression is mediated by the H3K27me3 repressive mark. In the second study, the Jmj16 H3K4me3 demethylase acts to keep SAGs repressed in younger leaves (Liu et al., 2019). In *jmj16* mutant alleles, both *WRKY53* and *SAG201* were up-regulated and associated with higher levels of the H3K4me3 mark. Non-catalytic forms of JMJ16 could bind to the promoter region, but only catalytically active forms could repress *WRKY53* gene expression. This second study demonstrated that changes in H3K4me3 marks can regulate SAGs.

*hac1* mutant alleles were reported to have darker green leaves (Li et al., 2014a). *HAC1* encodes a histone acetyl transferase from the CREB Binding Protein family (Bordoli et al., 2001; Pandey et al., 2001), which is known to acetylate histone H3 resulting in H3K9ac (Earley et al., 2007; An et al., 2017). H3K9ac is associated with open chromatin and increased gene expression, and genes directly regulated by HAC1 are expected to be down-regulated in *hac1* mutants. *hac1* mutants are pleiotropic and display a protruding gynoecium (Han et al., 2007). HAC1 also regulates flowering, and *hac1* mutants flower late due to increased *Flowering Locus C (FLC)* expression (Deng et al., 2007). FLC inhibits flowering, however decreased expression of genes that negatively regulate *FLC* was not observed in *hac1* mutants. HAC1 may have other non-histone targets or an unknown negative regulator of *FLC* could be down-regulated in late-flowering *hac1* mutants. In addition, *hac1/hac5* double mutant seedlings are hypersensitive to ethylene (Li et al., 2014b) and display the triple response (short root, short and thick hypocotyl and exaggerated apical hook) when grown in the dark in the absence of ACC, the non-gaseous precursor to ethylene. Neither single *(hac1* or *hac5)* mutant displayed ethylene hypersensitivity.

HAC1 also plays a role in the response to jasmonoyl-isoleucine (JA-ile), the active form of JA. HAC1 acetylates histones associated with MYC2 target genes to promote their expression. The Mediator Complex subunit, MED25 interacts with MYC2 and directly binds to and recruits HAC1 to target genes (An et al., 2017). Transcriptome data showed that genes induced by JA-ile were less responsive in a *hac1* mutant. In addition, genes co-regulated by JA-ile and HAC1 were enriched for many defense-related biological process GO terms as well as leaf senescence.

Here we show that *hac1* mutants have delayed age-related developmental leaf senescence. Potential HAC1 targets are identified by RNA-seq and ChIP-seq utilizing WT and two *hac1* alleles. T-DNA insertion mutants in three potential HAC1 targets were tested for leaf senescence phenotypes, and an *erf022* mutant disrupting the expression of *ERF022* showed delayed senescence. These findings implicate this AP2/ERF transcription factor as a novel positive effector of leaf senescence regulated by histone acetylation co-mediated by HAC1 and MED25.

## Materials and Methods

### Plant Growth Conditions

*Arabidopsis thaliana* Col-0 ecotype plants were grow in Sunshine^®^ Mix #1 Fafard^®^-1P RSi (Sungro Horticulture). The soil was treated with Gnatrol WDG (Valent Professional Products) (0.3 g/500 ml H2O) to inhibit the growth of fungus gnat larvae, and plants were sub-irrigated with Gro-Power 4-8-2 (Gro-Power, Inc.) (10 ml per gallon). Plants were grown in Percival AR66L2X growth chambers under a 20:4 light:dark diurnal cycle with a light intensity of 28 umoles photons m^-2^ sec^-1^. The low light intensity prevents light stress in older leaves, which was evident as anthocyanin accumulation at higher light intensities. To compensate for the reduced light intensity, the day length was extended. Leaves were marked by tying threads around the petioles soon after emergence from the meristem. Flowering time was determined when plants had 1 cm inflorescences (bolts). Leaf #5 from three week old plants were used for dark-induced senescence, and floated on water in the dark for the indicated number of days.

### Genotype analysis

Genomic DNA was isolated from two-three leaves using Plant DNAzol Reagent (ThermoFisher) following manufacturer’s instructions. Pellets were dried at room temperature for at least two hours, and resuspended in 30 uL TE (10 mM Tris, pH 8.0, 1 mM EDTA) overmight at 4°C. One microliter of genomic DNA was used as a template in PCR reactions with primers listed in Supplemental Table 2. All standard PCR reactions were performed with a 57°C annealing temperature using *Taq* polymerase with Standard *Taq* Buffer (New England Biolabs).

### Chlorophyll

One hole-punch was removed from each marked or detached leaf, and incubated in 800 μL N,N-dimethyl formamide (DMF) overnight in the dark. 200 μL of sample was placed in a quartz microplate (Molecular Devices) and readings were performed at 664 nm and 647 nm using a BioTek Synergy H1 plate reader. Absorbance readings were used to determine chlorophyll concentration (Porra et al., 1989). Chlorophyll was normalized to equal leaf area. For each genotype/condition, n =6.

### Total Protein

One leaf hole-punch was ground in liquid nitrogen in a 1.5 ml microfuge tube using a blue plastic pestle. 100 μL 0.1 M NaOH was added and the sample was ground for another 30 sec (Jones et al., 1989). Samples were incubated at room temperature for 30 min, centrifuged at 14000 rpm for 5 min. The Bradford protein assay (Bio-Rad Protein Assay Dye Reagent) was used to determine protein concentration in each supernatant using a bovine serum albumin standard. For each genotype/condition, n = 6.

### Percent Nitrogen

Elemental analysis for % nitrogen was done by Midwest Microlab, Indianapolis, IN. 100 dried seeds from one individual plant were in each sample (n = 8 for each genotype).

### Gene Expression

Total RNA was isolated from the Indicated leaves using Trizol reagent. 1000 ng of extracted RNA was used as a template for cDNA synthesis using MMLV-reverse transcriptase (New England Biolabs) and random hexamers to prime cDNA synthesis. The cDNA was diluted 16-fold and used as a template for real-time qPCR using either ABsolute QPCR Mix, SYBR Green, ROX (Thermo Scientific) or qPCRBIO SyGreen Blue Mix Hi-Rox (PCR Biosystems), in Step One Plus or Quant Studio 6 Flex qPCR machines. All real-time qPCR reactions used a 61°C annealing temperature.

For chlorophyll, total protein, percent nitrogen and gene expression, significant differences were determined using a t-test.

### RNA-seq

Indicated leaves were harvested and stored in liquid nitrogen. RNA was extracted and RNA-seq library production was performed using the breath adapter directional sequencing (BrAD-seq) method (Townsley et al., 2015). Real-time qPCR using ACT2 primers was the initial quality test. Libraries were sequenced at the Genome High-Throughput Facility (GHTF) at University of California, Irvine (UCI).

### ChIP-seq

Nuclei preparation and ChIP was performed as described previously (Brusslan et al., 2012). Libraries were produced and sequenced at the GHTF at UCI.

### Bioinformatics

RNA-seq raw data reads were aligned to the Arabidopsis TAIR 10 genome using Rsubread (Liao et al., 2013), and subject to quality control of count data and differential expression using NOISeq (Tarazona et al., 2015). The values were FPKM normalized using Tmisc and HTSFilter removed genes with low expression levels (Rau et al., 2013). A threshold value of q = 0.8 and a 2-fold change as the cut-off point was used to determine DEGs. ChIP-seq data were analyzed by MACS (Zhang et al., 2008) to find peaks of enrichment in comparison to input samples. MANorm (Shao et al., 2012) identified regions of differential histone modification. TopGO performed GO Biological Process enrichment and GAGE (Luo et al., 2009) performed pathway enrichment.

## Results and Discussion

### *hac1* Mutants Show Delayed Senescence

Two *Arabidopsis hac1* alleles *[hac1-1* (SALK_080380) and *hac1-2* (SALK_136314), Supplemental Figure 1] displayed darker green leaves when compared to WT. Age-related chlorophyll loss is shown in Figure 1A. At 28 days, total chlorophyll levels in leaf 7 were equal, but as the leaves aged, chlorophyll levels decreased faster in WT than the two *hac1* alleles. A significant difference in chlorophyll levels was detected between WT and both *hac1* alleles at day 48. The retention of chlorophyll was accompanied by reduced mRNA levels for genes associated with leaf senescence (Figure 1B). *AtNAP* encodes a positive regulator of leaf senescence associated with ABA synthesis (Liang et al., 2014; Yang et al., 2014). *NIT2* encodes a nitrilase that is highly expressed in leaf senescence, and contributes to auxin synthesis (Normanly et al., 2007) and glucosinolate catabolism (Vorwerk et al., 2001). *NYC1* encodes a chlorophyll *b* reductase required for light harvesting complex disassembly (Kusaba et al., 2007). The chlorophyll and gene expression data show that *hac1* alleles display delayed leaf senescence.

**Figure 1.**
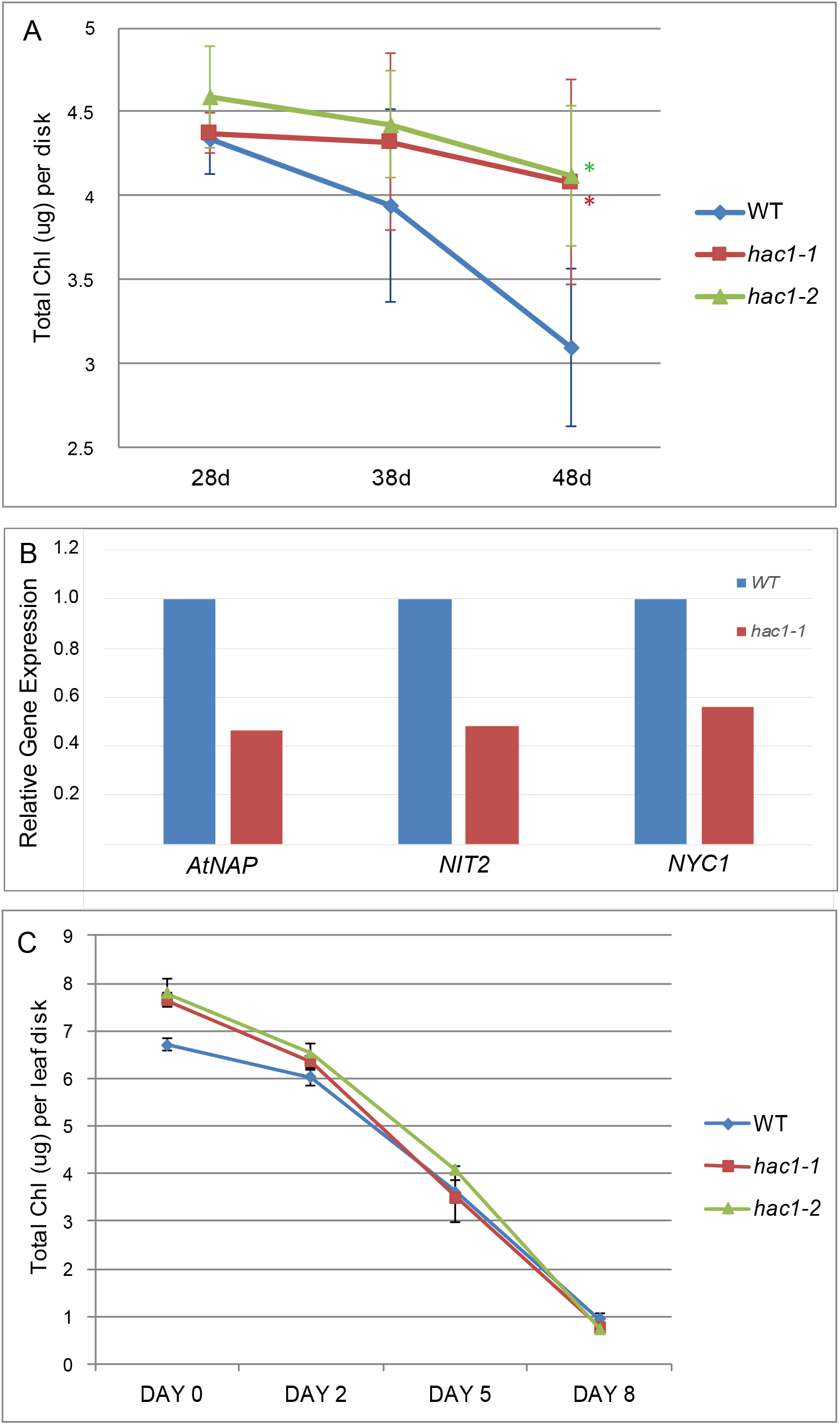
Delayed age-related senescence in *hac1* alleles. A) To observe age-related developmental senescence, total chlorophyll was measured from leaf 7 from plants that had grown 28, 38 or 48 days. Significant differences from WT are indicated by asterisks (t-test, p < 0.05) and were observed for both *hac1* alleles at 48 days. B) RNA was extracted from WT and the *hac1-1* allele at 48 days from leaf 6 of the same plants shown in panel A, and gene expression for three SAGs was measured by real-time qPCR. C) Leaf 5 was removed from plants grown for 21 days, and floated on water in the dark for the indicated number of days to observe dark-induced senescence. One leaf disc was removed from each leaf and chlorophyll was measured. No significant differences were observed. All error bars show the 95% confidence interval.

The reduction of total chlorophyll was also evaluated in detached leaves floated in water in the dark (dark-induced senescence), and no difference was noted between WT and the two *hac1* alleles (Figure 1C). There are molecular differences in the signaling pathways between dark-induced and developmental senescence; most prominent is the role of SA in developmental, but not in dark-induced senescence (Buchanan-Wollaston et al., 2005; van der Graaff et al., 2006; Guo and Gan, 2012). Thus, it is possible that alterations in the signaling of developmental senescence do not necessarily accompany changes in dark-induced senescence. These results support a role for HAC1 as a promoter of age-related, developmental leaf senescence.

A trending increase in total leaf protein concentration accompanied the significant increase in chlorophyll levels in both *hac1* alleles (Figures 2A-B). However, the delayed senescence in the *hac1* alleles did not result in greater percentage of seed nitrogen (Figure 2C). Delayed senescence in wheat was reported to increase grain nitrogen concentration (Zhao et al., 2015), however the relationship between percentage of seed nitrogen and leaf senescence is complex (Chardon et al., 2014; Havé et al., 2017).

**Figure 2.**
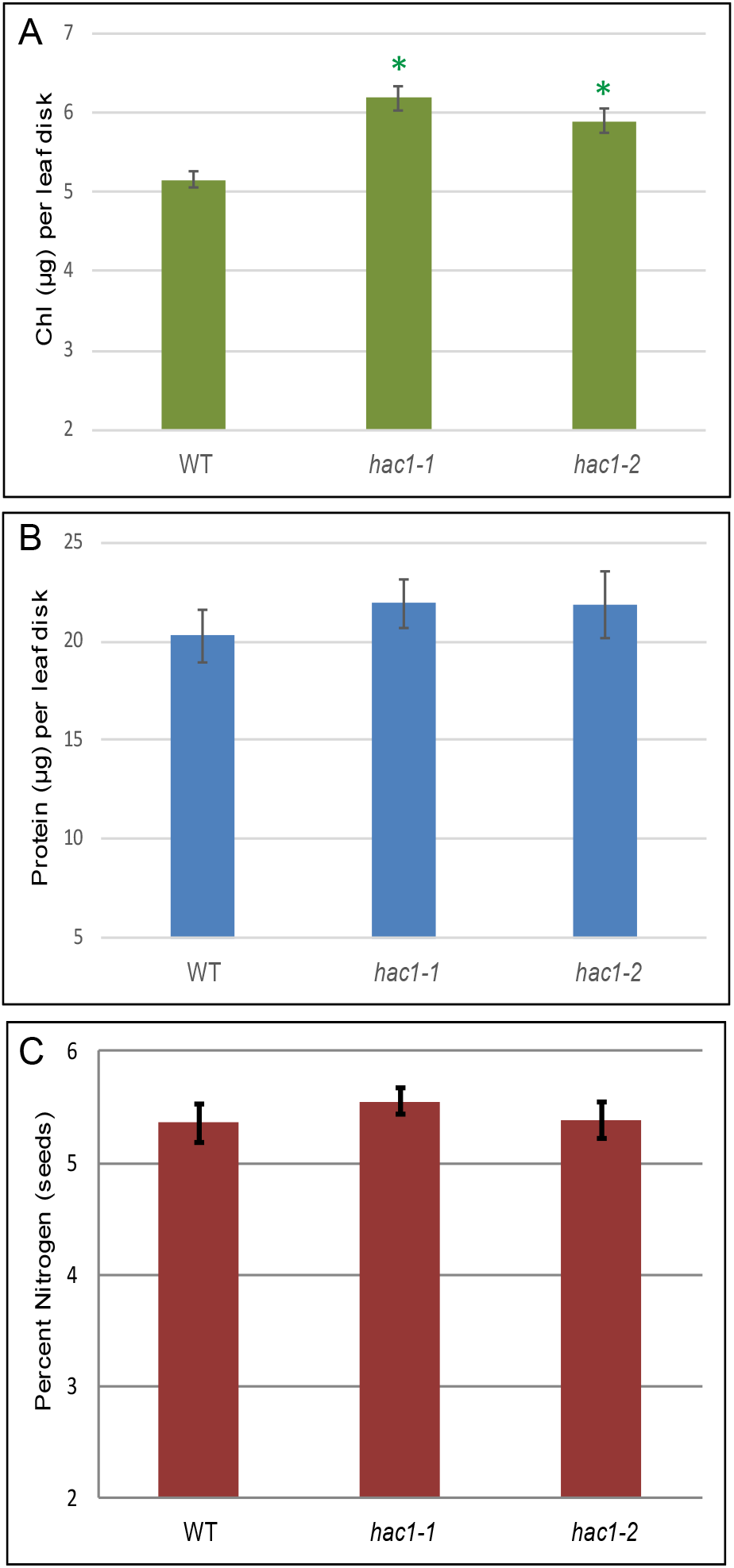
Chlorophyll, protein and seed nitrogen content in *hac1* alleles. Plants were grown for 49 days and chlorophyll (A) and total protein (B) were measured in hole-punch disks from leaves 12-14, n = 8. Significant differences between WT were observed for chlorophyll, but not total protein (t-test, p < 0.05). Seeds were harvested from individual plants and batches of 100 dried seeds were subject to elemental analysis (C). No significant differences in percent nitrogen were observed, n = 8. All error bars show the 95% confidence interval.

### *hac1* Mutants Display Altered Levels of Histone Modifications and Changes in Gene Expression During Leaf Senescence

ChIP-seq was performed on the same tissue shown in Figure 2 to identify genes associated with a loss of H3K9ac and/or H3K4me3 histone modifications in both *hac1* alleles. HAC1 catalyzes H3K9 acetylation, and both H3K9ac and H3K4me3 are associated with active gene expression (Berr et al., 2011). As expected, H3K9ac significantly decreased at 968 loci and increased at only 555 loci in both *hac1* alleles. H3K4me3 modifications were similarly affected, with 548 loci showing a loss and only 33 loci showing a gain of H3K4me3 marks. RNA-seq was used to identify differentially expressed genes (DEGs) between WT and both *hac1* alleles. Accordingly, the number of up-regulated DEGs (12) was much smaller than the number of down-regulated DEGs (143) in both *hac1* alleles. These 143 down-regulated DEGs were subject to pathway enrichment analysis, and significant enrichment of glucosinolate biosynthesis, plant-pathogen interaction, as well as glutathione and ascorbic acid metabolism were revealed. These pathways are stress-related and their down-regulation in *hac1* likely slows the rate of leaf senescence. One GO term enriched in the up-regulated DEGs in both *hac1* alleles is ribosome biogenesis, which occurs during rapid protein synthesis, and would be important for anabolic growth, not catabolic senescence. Cytokinin action delays dark-induced senescence, in part, by maintaining the expression of genes associated with ribosome GO terms (Kim et al., 2018b).

The Venn diagram in Figure 3 shows the overlap of genes with reductions in H3K9ac and H3K4me3 marks, as well as decreased expression in both *hac1* alleles. Our analysis identified 44 genes (Supplemental Table 2) with reductions in H3K9ac marks and gene expression. These potential HAC1 targets have enriched GO terms including response to chitin and response to abiotic stimulus. These GO biological process terms have previously been associated with SAGs (Brusslan et al., 2015). Two of the potential HAC1 targets, *IGMT1* and *CYP81F2* (green highlight in Supplemental Table 2), encode indole glucosinolate biosynthetic enzymes, providing evidence that these secondary compounds are important during leaf senescence and potentially regulated via histone acetylation. We also observed significant reductions in H3K4me3 marks for these two genes in both *hac1* alleles, further bolstering the presence of chromatin changes.

**Figure 3:**
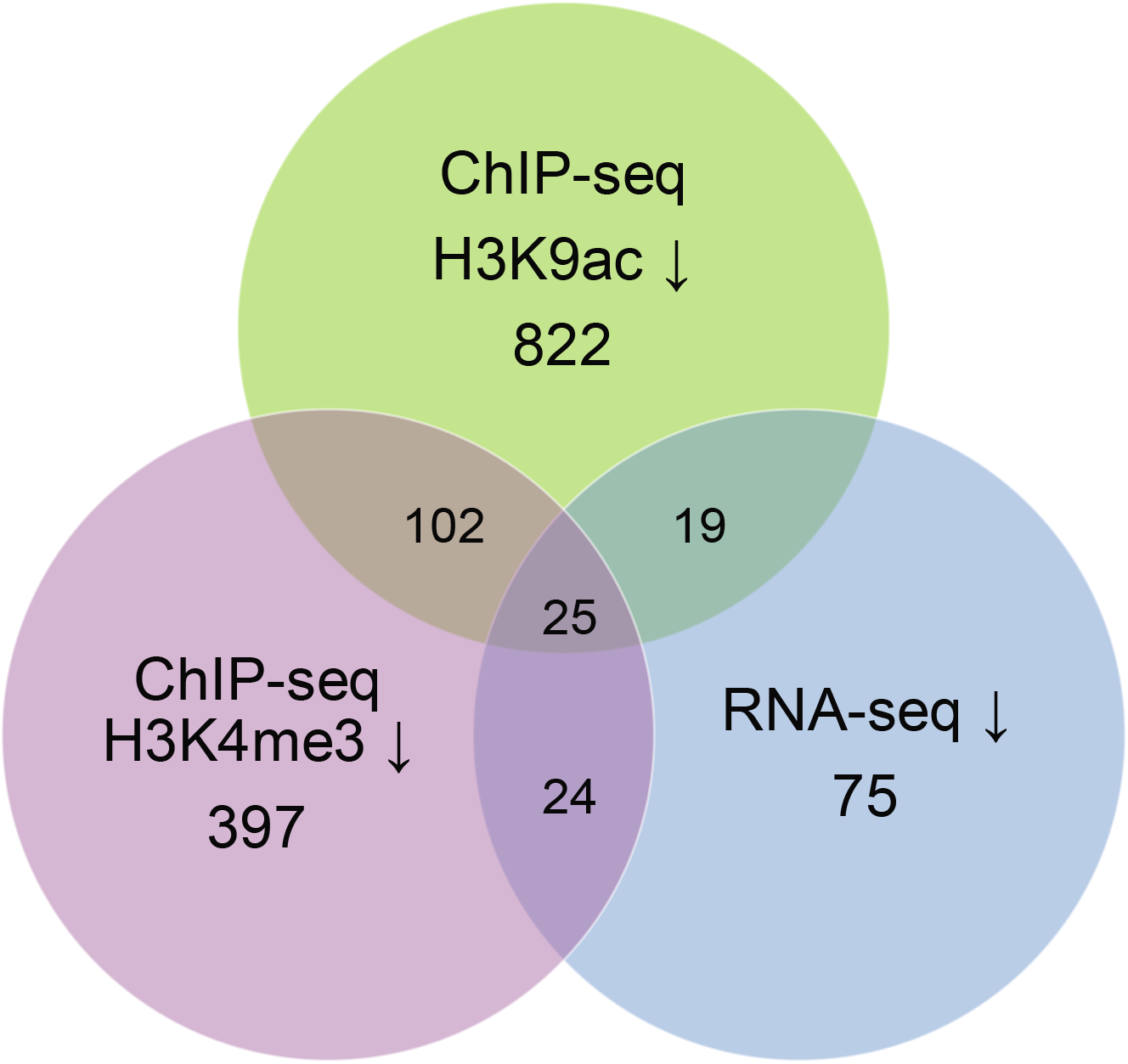
The Venn diagram shows the overlap of genes with reductions in gene expression and histone modifications. WT, *hac1-1* and *hac1-2* (49 days, leaf 12-14) were subject to RNA-seq and ChIP-seq using H3K9ac and H3K3me3 antibodies. Genes that showed a significant reduction in both *hac1* alleles in comparison to WT were considered to have lower expression (RNA-seq) or reduced histone marks (ChIP-seq).

### Analysis of Leaf Senescence Phenotypes in Potential HAC1 Targets

We measured leaf senescence in T-DNA insertion lines disrupting three regulatory genes from the list of 44 potential HAC1 targets (yellow highlights in Supplemental Table 2). These include *ERF022, MYB15* and *TMAC2.* Two of these genes: *ERF022* and *TMAC2* also show a reduction in H3K4me3 marks. *ERF022* and *MYB15* encode transcription factors while TMAC2 plays a negative role in ABA response (Huang and Wu, 2007). Flowering time, *NIT2* gene expression, and chlorophyll levels were quantified in these mutants (Figure 4A-C). We also showed that full-length mRNAs spanning the T-DNA insertion were not produced in each mutant allele (Figure 4D). The only line to show a consistent and strong significant alteration in leaf senescence was *erf022*, with slightly later flowering (by about three days), and after 44d of growth, reduced *NIT2* expression (approximately 8-fold) and increased chlorophyll. These phenotypes indicate a delay in leaf senescence and implicate *ERF022* as a positive regulator of leaf senescence.

**Figure 4:**
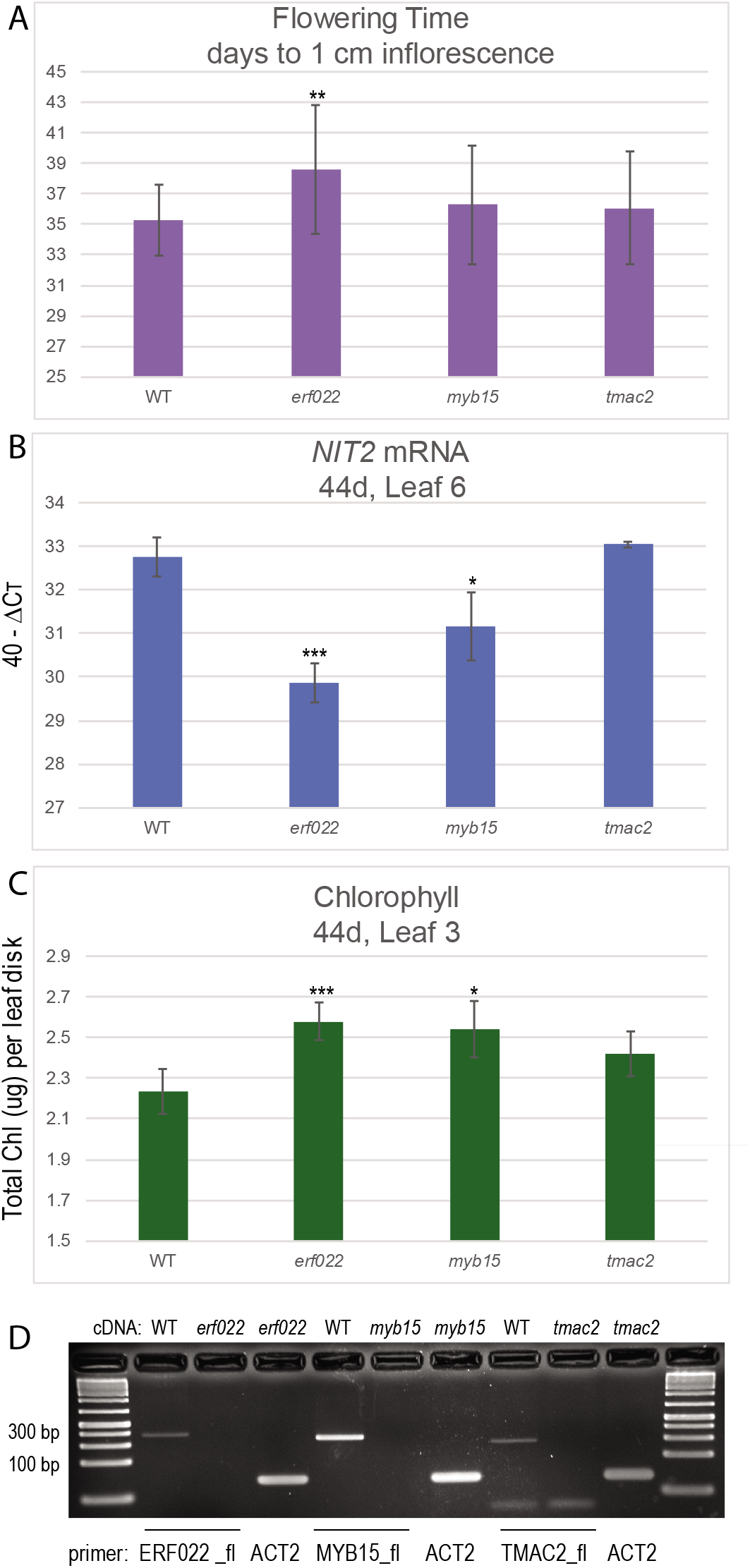
Senescence phenotypes in T-DNA insertion lines disrupting potential HAC1 target genes. Panel A shows flowering time and error bars show the standard deviation of two separate trials. Panel B shows *NIT2* gene expression and panel C shows total chlorophyll (n =6 for all genotypes). One biological replicate is shown, however similar results were obtained in a second biological replicate. Error bars for panels B and C show the 95% confidence interval. A t-test was used to evaluate significant differences: * = p<0.05, ** = p<0.01, *** = p<0.001. Gene expression is measured as 40 – ΔCt. The ΔCt value is the Ct value of ACT2 – the Ct value of the gene of interest. Panel D shows that full-length mRNAs were not produced in T-DNA insertion alleles. The cDNAs templates are shown above the PCR products and the primers are shown below. Primer sequences are available in Supplemental Table 1.

Our results suggest that H3K9 acetylation mediated by HAC1 occurs at *ERF022* during leaf aging, and is accompanied by changes in H3K4me3 marks. Together, these two marks likely promote the expression of *ERF022*, a positive regulator of leaf senescence. ERF022 is a member of the drought-responsive element-binding (DREB) subfamily of the AP2/ERF family (Nakano et al., 2006). Protoplast transfection experiments show ERF022 to be a positive regulator of the RD29A promoter (Wehner et al., 2011), suggesting ERF022 may mediate abiotic stress. Etiolated *erf022* mutant seedlings produce significantly more ethylene, suggesting that ERF022 attenuates ethylene synthesis early in development (Nowak et al., 2015). *EIN2* encodes an essential component of the ethylene signaling pathway, and *ein2* mutants delay leaf senescence (Oh et al., 1997), thus increased ethylene production would be expected to accelerate senescence. If ERF022 is acting similarly in seedlings and older leaves, increased ethylene would be expected to promote senescence, however a delay was observed in *erf022.* It is possible that ERF022 plays different roles at different times in development. JA and a necrotrophic pathogen stimulated *ERF022* expression (Mcgrath et al., 2005), indicating ERF022 plays a role in defense. Defense and senescence share many genes, as noted previously. Of interest, the ethylene hypersensitivity previously observed in *hac1/hac5* double mutant seedlings may be due to reduced expression of *ERF022. erf022* mutants overproduce ethylene, and mutations in HAC1 and HAC5 additively displayed a constitutive triple response.

### MEDIATOR25 works additively with HAC1 to regulate *ERF022* expression

The MED25 subunit of the Mediator Complex can interact with HAC1. We obtained *med25* mutants and produced *hac1-1/med25* double mutants to evaluate genetic interaction. The longest delay in flowering was observed for *med25* and *hac1-1/med25* (Figure 5A), but an additive effect in flowering phenotype was not present. Chlorophyll levels were measured in leaf 7 in 45 day old plants, and higher chlorophyll levels were observed in *hac1-1, med25* and the *hac1-1/med25* double mutants, and although all lines were significantly greater than WT, none were significantly different from each other (Figure 5B). These data suggest that HAC1 and MED25 do not have an additive effect, as loss of one or both show similar delays in flowering and chlorophyll loss. The *erf022* mutant was also included in this experiment; it bolted later and had more chlorophyll than WT, but it did not differ from the *hac1-1, med25* or *hac1-1/med25* mutant lines.

**Figure 5:**
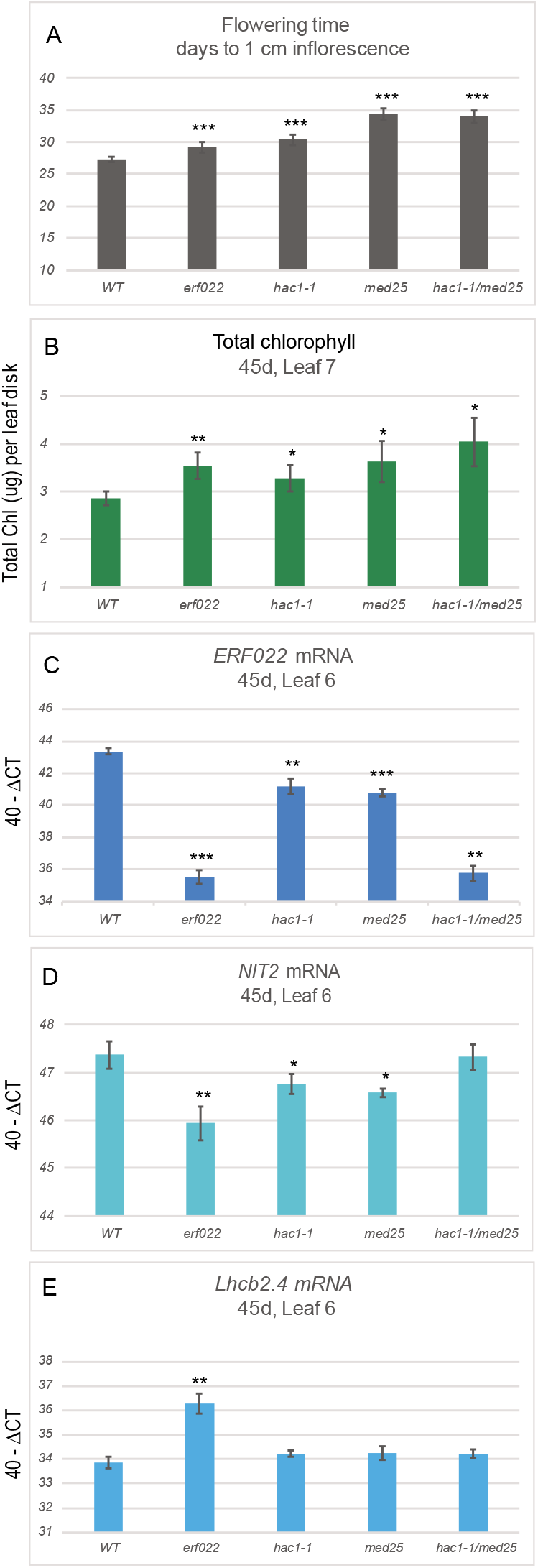
Senescence phenotypes in *hac1-1/med25* double mutants. All lines were evaluated for flowering time (panel A). At 45 days of growth, chlorophyll was measured in leaf 7 and RNA was extracted from leaf 6. Total chlorophyll levels (μg per leaf disk) are shown in panel B. *ERF022* (panel C), *NIT2* (panel D) and *Lhcb2.4* (panel E) mRNA levels are shown. A t-test was used to evaluate significant differences: * = p<0.05, ** = p<0.01, *** = p<0.001. All error bars show the 95% confidence interval, n=6 for all genotypes.)

Gene expression was also evaluated in these mutant lines. As expected, *ERF022* expression was minimally detected in the *erf022* mutant. A strong additive effect was seen between *hac1-1* and *med25* with much lower *ERF022* expression in the *hac1-1/med25* double mutant than in either single mutant (Figure 5C). These data suggest that MED25 guides HAC1 to histones at the *ERF022* locus to direct histone acetylation for increased chromatin accessibility. With respect to two other SAGs, *NIT2* and *Lhcb2.4*, the *erf022* mutant showed the largest effect: minimal up-regulation of *NIT2* (Figure 5D) and minimal down-regulation of *Lhcb2.4* (Figure 5E) as compared to *hac1-1, med25* and *hac1-1/med25.* These data suggest that loss of ERF022 has a more profound effect on the leaf senescence phenotype than its down-regulation through loss of both HAC1 and MED25. Although the *ERF022* transcript levels were similar to the *hac1-1/med25* double mutant (Figure 5C), it is probable that the mRNA produced in the *erf022* mutant is inefficiently translated due to the T-DNA insertion in the 3’-UTR and led to a stronger phenotype in *erf022.* In addition, there are likely more genes mis-regulated in *hac1-1/med25* and these may have compensating effects on leaf senescence.

## Conclusion

*hac1* mutant alleles display a delay in leaf senescence implicating histone acetylation as a contributor to the regulation of leaf senescence. A combined approach using ChIP-seq, RNA-seq and genetic analysis, identified ERF022 as a novel positive effector of leaf senescence regulated by H3K9ac and H3K4me3 marks. *ERF022* is possibly a direct target of HAC1, which operates in concert with MED25 to allow full expression of *ERF022* in older leaves.

## Supporting information

Hinckley_etal_Supplemental

## Acknowledgements

The authors thank Soumi Barman and Glenn Nurwano for technical help in genotype analysis. Research reported in this publication was supported by the National Institute of General Medical Sciences of the National Institutes of Health under Award Numbers R25GM071638 and SC3GM113810. The content is solely the responsibility of the authors and does not necessarily represent the official views of the National Institutes of Health.

## Author Contributions

WEH and KK designed and performed the research and analyzed data. JAC performed the research. JAB designed and performed the research, analyzed data and wrote the paper. All authors greatly contributed to editing.

## Supplemental Data files

Supplemental Figure 1: Full-length mRNAs are not produced in *hac1* alleles.

Supplemental Table 1: Primers

Supplemental Table 2: Genes with decreased H3K9ac and mRNA in both *hac1* alleles RNA-seq and ChIP-seq data files are in the process of being added to the NCBI GEO database.

