## Supplementary material for "The *HAC1* Histone Acetyltransferase Promotes Leaf Senescence via Regulation of *ERF022*": Hinckley_etal_Supplemental

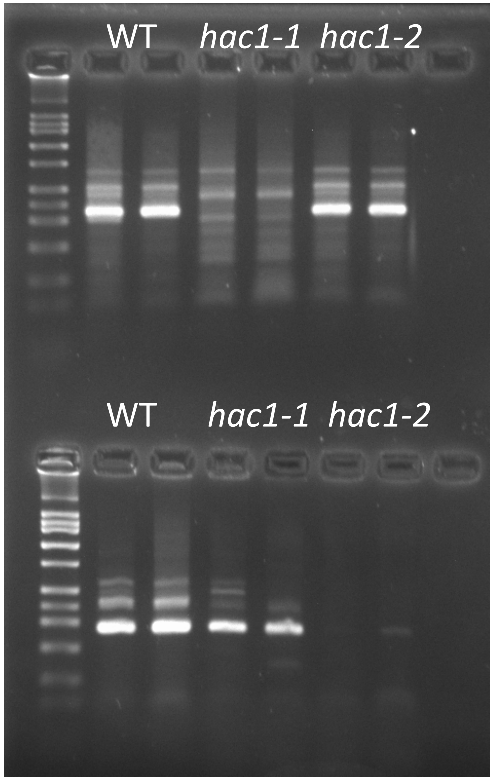


Supplemental Figure 1. No full-length transcripts are produced in *hac1* alleles.

Top: SALK_080380 (*hac1-1*) harbors a T-DNA insertion in exon 6 and primers flanking this exon do not amplify *hac1-1* cDNA.

Lower: SALK_136314 (*hac1-2*) harbors a T-DNA insertion in exon 12 and primers flanking this exon do not amplify *hac1-2* cDNA.

The exon 12 primers do amplify *hac1-1* cDNA indicating that a partial transcript downstream from the T-DNA is produced. The exon 6 primers amplify *hac1-2* cDNA indicating that a transcript upstream of the T-DNA is produced. No full length *HAC1* transcripts are produced in either mutant allele.

### Supplemental Table 1: Primers

| Primers for amplification of genomic DNA to determine if lines homozygous for T-DNA insertions (57$^{o}$C annealing temperature) | | | T-DNA Insertion |
|---|---|---|---|
| Primer Name | Sequence (5' to 3') | | |
| LBb1.3_plus | GGATTTTGCCGATTTCGGAACC | | |
| _080380C_LP | AGGGAACATGTCATCCATGAG | *hac1-1* | SALK_080380 |
| _080380C_RP | AGGGAGAATCTGAGAACCTGC | | |
| 136314C_LP | TCAGTATGCAGCAATGAGTGC | *hac1-2* | SALK_136314 |
| 136314C_RP | TCGGGAGGATAGTTACCCAAC | | |
| SALK_091690_LP2 | AACCTTGGTTGATGATATGAC | *erf022* | SALK_091690 |
| SALK_091690_RP | GAACCCGGTAAGAAAACAAGG | | |
| SALK_151976_LP | GAAGATGGGGTTGAAGAGAGG | *myb15* | SALK_151976 |
| SALK_151976_RP | CTAAGAGATCTTGTTCCCCGC | | |
| SALK_208284_LP | TGTTGGTAGTCTTGGCTTTGC | *tmac2* | SALK_208284 |
| SALK_208284_RP | GGGTCGATCTTCTTCGATACC | | |
| SALK_129555_LP | TGGAACTGGTCCAACAGAAAC | *med25* | SALK_129555 |
| SALK_129555_RP | TGCATTGGCTTTCTTCCATAC | | |

**Primers for amplification of cDNA to determine if mRNA spanning T-DNA insertion site is produced (57$^{o}$C annealing temperature)**

| | | |
|---|---|---|
| 080380C_Fw | TAAATCTCTGTTGCTGCTGGC | *hac1-1* |
| 080380C_Rev | GATTCAGGTACATCCTACACC | |
| 136314C_Fw | AGCTCTACATTCACCAGTTTGC | *hac1-2* |
| 136314C_Rev | AACGGAGAAGCTCTGCGTAC | |
| ERF022F_fl | TACAGAGGCATTCGTCGGA | *erf022* |
| ERF022R_fl | TCGGCGAGTCTAATGGAGAC | |
| MYB15F_fl | TAGAAGCGAATCGGAGCTAG | *myb15* |
| MYB15R_fl | CAGCCACTTCTAGGTCATTC | |
| TMAC2F_fl | ACACACAAGGCATGTCTTCA | *tmac2* |
| TMAC2R_fl | GTCGTAACGGAGATGGATAT | |
| kfMed25F_fl | GCCAACACCTGTATATCGTC | *med25* |
| kfMed25R_fl | TGGGCTTTCCCGCATTGTAT | |

**Primers for real-time qPCR (61$^{o}$C annealing temperature)**

| | |
|---|---|
| ACT2_F | GCGACTTGACAGAGAAGAAC |
| ACT2_R | GAAAGAGCGGAAGAAGATGAG |
| NIT2_F | TACAACGATACTCCCGCCAC |
| NIT2_R | CACCCCTAAACCAAACCTAAAAC |
| AtNAP_F | CTTCCCAATCTACCCTCCCT |
| AtNAP_R | GGAGACAGGGCATGGTTTAG |
| NYC1_F | ATGGGTCGTGGGATTCTTTG |
| NYC_R | GCAATTTGAACAGCAACCGA |
| erf022qp2F | TCTCCATTAGACTCGCCGAG |
| erf022qp2R | AAACTCATCCCTATCGTGCTGA |
| Lhcb2.4_F | CCGAAAACACACCATCATACC |
| Lhcb2.4_R | TCCCAACATTGCCCATCTAC |

**Supplemental Table 2: Decreased H3K9ac and mRNA in both *hac1* alleles**

| | |
|---|---|
| AT1G31580 | ECS1, cell wall |
| AT1G33760 | ERF022 |
| AT1G36622 | transmembrane |
| AT1G56250 | PP2-B14, phloem, F-box E3 ubiquitin ligase |
| AT1G61260 | DUF761 |
| AT1G66180 | putative aspartyl protease |
| AT1G67810 | SUFE2, chloroplast Fe-S cluster assembly |
| AT1G02400 | GA2OX6, GA oxidase |
| AT1G15830 | hypothetical |
| AT1G21100 | IGMT1, indole glucosinolate O-methyltransferase |
| AT1G24575 | DEAD-box ATP-dependent RNA helicase-like protein |
| AT1G27020 | plant/protein |
| AT2G32030 | Acyl-CoA N-acyltransferases (NAT) superfamily |
| AT2G35930 | PUB23, U-Box E3 ubiquitin ligase |
| AT2G38500 | 2-oxoglutarate (2OG) and Fe(II)-dependent oxygenase superfamily |
| AT2G41100 | TCH3, calmodulin-like |
| AT2G15090 | KCS8, very long chain fatty acid biosynthesis |
| AT2G22860 | PSK2, phytosulfokine 2 peptide precursor |
| AT3G29670 | PMAT2, phenolic glucoside malonyltransferase |
| AT3G48520 | CYP94B3, catalyzes the formation of 12-OH-JA-Ile from JA-Ile: attenuates JA signaling. |
| AT3G50930 | BCS1, mitochonrial outer wall, BC1 synthesis, amplifies SA signaling |
| AT3G52450 | PUB22, U-Box E3 ubiquitin ligase |
| AT3G01830 | Calcium-binding EF-hand family |
| AT3G10985 | SAG20, response to Nep1 (necrotic fungal protein) |
| AT3G02140 | TMAC2, negative regulator of ABA |
| AT3G16530 | Lectin like |
| AT3G23250 | MYB15, basal immunity, Putative WRKY53 target |
| AT4G27300 | S-locus lectin protein kinase |
| AT4G30280 | XTH18, xyloglucan endotransglucosylase/hydrolase |
| AT4G36950 | MAPKKK21, mitogen activated protein kinase kinase kinase |
| AT4G37610 | BT5, BTB and TAZ domain protein 5, high redundancy within 5 member family |
| AT4G01360 | BPS3, BYPASS 3, Down-regulate BYPASS active mobile compound |
| AT4G15760 | MO1, monooxygenase 1, similar to SA degradation enzymes |
| AT5G05600 | JAO2, JA oxidase 2 |
| AT5G38700 | cotton fiber |
| AT5G57220 | CYP81F2, indole glucosinolate biosynthesis, I3G to 4OH-I3G |
| AT5G60910 | AGL8, FRUITFULL, MADS-box |
| AT5G62165 | AGL42, FOREVER YOUNG FLOWER, Delays flower senescence |
| AT5G62360 | PMEI13, pectin methyl-esterase inhibitor 13, salt tolerance |
| AT5G62520 | SRO5, ADP ribosylation |
| AT5G66650 | calcium uniporter (DUF607) |
| AT5G10760 | AED1, aspartyl protease, systemic acquired resistance |
| AT5G13100 | Gap junction beta-4 |
| AT5G19230 | Glycoprotein membrane precursor GPI-anchored |
